# Developmental timeline of the lateral olfactory tract innervation of piriform cortex

**DOI:** 10.64898/2026.11.24.768942

**Authors:** Eduardo Martin-Lopez, Haolin Li, Natalie J. Spence, Nathan Bauman, Jose M. Estrada-Giron, Bowen Brennan, Christine Kaliszewski, Charles A. Greer

## Abstract

The development of the lateral olfactory tract (LOT) begins embryonically and is closely linked to the differentiation of olfactory bulb (OB) projection neurons, mitral and tufted cells, whose axons form the LOT. Here, we implemented calretinin (CR) as a proxy to characterize LOT development. Immunohistochemistry (IHC) for CR revealed expression in most mitral/tufted cells, with lower levels in the dorsal OB. This mirrored a reduced CR labeling in a restricted ventral LOT domain, suggesting topographic organization of OB projections. To achieve axonal resolution, we used Calb2-CreERT2 × Ai14 mice to label CR^+^ neurons and their axons with tdTomato. Expression began at E12, when pioneer LOT axons reach the anterior piriform cortex (APC) and posterior brain regions. From E13-E14, axon density increased, contributing to LOT thickening and extension toward the amygdala. Axonal arborization was not evident in the posterior piriform cortex (PPC) until E15. Electron microscopy revealed excitatory asymmetric and inhibitory symmetric synapses in the APC as early as E13, with relatively stable embryonic densities. Postnatally, excitatory synapses increased in both APC and PPC between P9 and P21. Collectively, this work establishes a developmental timeline of LOT formation and sheds light on fundamental aspects of the assembly of olfactory cortical circuits.

## Introduction

The primary olfactory system is an ancient and discriminative chemosensory system that detects airborne chemicals (odorants) in the environment. Beginning in the olfactory epithelium (OE), a sensory organ located in the nasal cavity (Firestein, 2001), the olfactory pathway follows two sequential relays. First, the olfactory sensory neurons (OSNs) in the OE detect and transduce odor signals into action potentials that are sent to the olfactory bulb (OB) via the OSN axons that form the olfactory nerve (ON). In the OB the OSN axons coalesce into odorant receptor (OR) specific glomeruli (Mombaerts et al., 1996; Treloar et al., 2002). Second, following odorant specific initial processing in the OB, the projection neurons, mitral and tufted cells, transmit the olfactory information to the olfactory cortex (piriform cortex, PC), ventral striatum (olfactory tubercle, OT) and limbic regions such as the amygdala (AMY) and entorhinal cortex (EC) via the lateral olfactory tract (LOT) (Powell et al., 1965; Igarashi et al., 2012; Shepherd et al., 2021). In rodents, LOT axons run parallel to the rostral-caudal axis of the brain in the ventrolateral region of the frontal and parieto-temporal lobes, where the olfactory information is distributed to broad ensembles of neurons for odor processing and representation (Stettler and Axel, 2009; Ghosh et al., 2011; Bekkers and Suzuki, 2013; Imamura et al., 2020; Zeppilli et al., 2021; Chae et al., 2022; Chen et al., 2022; Berners-Lee et al., 2023; Stenwall et al., 2025).

Developmentally, the first sign of OSNs differentiation occurs as early as the embryonic day 9 (E9) in the mouse olfactory placode with the first evidence of OR expression (Miller et al., 2010b; Rodriguez-Gil et al., 2010; Tufo et al., 2022). These nascent OSNs extend pioneer axons by E10 which reach the OB one day later at E11 in the mouse (Marin-Padilla and Amieva, 1989; De Carlos et al., 1996; Whitesides and LaMantia, 1996; Rodriguez-Gil and Greer, 2008; Shay et al., 2008; de Castro, 2009; Miller et al., 2010a; Rodriguez-Gil et al., 2015). In mice, the first evidence of mitral cells exiting the cell-cycle occurs at E9 (Imamura et al., 2011; Imamura and Greer, 2015), but it is not until E11-12 that the first ones are identified as migratory bipolar neurons oriented tangentially to the surface of the presumptive OB (Hinds, 1968b; 1972; Lopez-Mascaraque and de Castro, 2002; Blanchart et al., 2006; Miller et al., 2010a). This raises two fundamental questions: when do the first LOT axons emerge, and how do they extend during embryonic development?

Analyses of OB embryogenesis in mice using Golgi impregnations suggested that the first mitral cells axons are detected at E13 (Hinds, 1972). Subsequently, Sato et al. (1998) used a fluorescent anterograde dextran and reported that the first LOT axons appeared at E12, confined within the primordium of the OB (Sato et al., 1998). In 2006, Walz and colleagues developed GFP in neurotensin^+^ as a proxy for mitral/tufted cell axons and reported innervation within the LOT in the primordium of the OB at E11.5 (Walz et al., 2006). In parallel, whole-mount immunohistochemistry against L1 and neuropilin-1/2, markers of mitral/tufted cells and their axons, revealed that LOT axons are detectable at E12 and have extended across the entire PC at this stage (Inaki et al., 2004). From E13 onwards it seems clear that LOT axons have reached the entire surface of PC, EC and AMY (Tomioka et al., 2000; Lopez-Mascaraque and de Castro, 2002; Inaki et al., 2004), supported by a conduit of LOT guidepost cells (Sato et al., 1998; Hirata et al., 2012). These discrepancies, likely related to the different techniques used to track the developing LOT axons, highlight the need to revisit LOT development using more contemporary approaches. Therefore, here we sought to shed new light on the developmental timeline of LOT axons in parallel with PC synaptogenesis. As a proxy to track the emergence and development of the mitral/tufted cell axons we implemented calretinin which is expressed prior to axon extension and is specific to the LOT axons.

Calretinin (CR), also known as Calbindin-2 (Calb2), is a calcium binding protein first described in the chick retina (Rogers, 1987). Its expression is characteristic of some GABAergic interneurons throughout the brain and regulates intracellular calcium signaling and neuronal excitability (Resibois and Rogers, 1992; Rogers and Resibois, 1992; Barinka and Druga, 2010; Cauli et al., 2014; Vanessa Becerra-Hernandez et al., 2025). In the adult olfactory system, CR primarily labels subpopulations of interneurons distributed throughout the OB, the anterior olfactory nucleus (AON), PC and intermingled among axons from the anterior commissure (AC) (Whitman and Greer, 2007; Gavrilovici et al., 2010; Kay and Brunjes, 2014; Martin-Lopez et al., 2018; Martin-Lopez et al., 2019a; Capsoni et al., 2021). However, in the OB CR is also expressed by mitral/tufted cells, the glutamatergic projection neurons of the OB, and consequently by their axons (Jacobowitz and Winsky, 1991; Resibois and Rogers, 1992; Wouterlood and Härtig, 1995; Pombal et al., 2002). This labeling makes CR an opportunistic marker to identify the LOT and boundaries between sublayers of PC (Sarma et al., 2011; Amaniti et al., 2015; Martin-Lopez et al., 2019a; Martin-Lopez et al., 2019b; Moreno-Velasquez et al., 2020).

Here, we initially used CR immunohistochemistry to characterize its expression in the OB and PC. In the PC, CR labeled the LOT and layer Ia, while in the OB, it was expressed by mitral/tufted cells throughout the entire circumference, with lower expression in the dorsal region. Of interest, this regional variation correlated with a reduced CR labeling in a restricted domain of LOT axons. Because CR immunolabeling did not provide sufficient axonal resolution, we used CR-CreERT2 mice (Taniguchi et al., 2011) crossed with the Ai14 reporter mice (Madisen et al., 2010) to label CR^+^ mitral/tufted cells and their axons with tdTomato. We then systematically examined LOT development from E12 onward to establish the timeline of axon production, elongation, and extension throughout PC. In addition, electron microscopy was used to assess LOT axon density and synaptogenesis in PC during development. Collectively, our analyses establish a timeline for mitral/tufted cell innervation of olfactory cortical regions via the LOT and identify CR as a useful marker for tracing LOT axons, providing a new approach to studying the development of higher olfactory cortical regions.

## Materials and Methods

### Animals

All experiments performed to test the expression and distribution of CR by the OB projection neurons, mitral/tufted cells, and those for EM analysis, were carried out on wild-type CD1 mice obtained from Charles River (cd-1r-igs) at adult and embryonic stages. To document the expression of CR by mitral/tufted cells, we used brain tissues from adult CD1 mice ordered at 8-10 weeks of age. To study the distribution of CR^+^ projection neurons along the OB cardinal axes, we used young adults at postnatal day 33 (P33) that were injected with AAVs at P0 in their transverse sinuses to label neuronal somas and axons. Therefore, for these experiments we ordered pregnant females at embryonic day 16 (E16) and maintained them in our facility until delivery date. For EM analysis, we ordered CD1 pregnant females at E11 and maintain them in our vivarium to be used at different ages as follows: one group were used embryonically to obtain tissues from E13 to E17, while a second group were maintained until delivery to obtain postnatal tissues at P1, P9 and P21 ages.

The experiments designed to track the development of the LOT into PC along the anterior-posterior axis were carried out using the endogenous expression of the fluorophore tdTomato in CR^+^ cells using a transgenic mice that we obtained by crossing the following two strains: 1) the B6(Cg)-Calb2^tm2.1(cre/ERT2)Zjh^/J mouse strain (The Jackson Laboratory, RRID:[IMSR_JAX:013730](https://www.jax.org/strain/013730)), also known as Calb2-CreERT2 or CR-CreERT2, has Calb2 (calbindin 2; also called CR) gene function abolished and contains direct CreERT2 fusion protein expression in neurons regulated by the Calb2 promoter/enhancer elements (Taniguchi et al., 2011); this strain was crossed with 2) the B6.Cg-Gt(ROSA)26Sor^tm14(CAG-tdTomato)Hze^/J reporter mice (The Jackson Laboratory, [RRID:IMSR_JAX:007914](https://www.jax.org/strain/007914)), also known as Ai14, that have robust tdTomato expression following Cre-mediated recombination (Madisen et al., 2010). The heterozygous offspring (named here as Calb2-CreERT2 × Ai14) guaranteed the expression of both CR and tdTomato from the Ai14 allele after injections of 4OH-Hydroxytamoxifen (4OHTx) (Fig. 3A). While some mice were used at adult stages (8-10 weeks) and injected with 4OHTx postnatally, others were used embryonically at consecutive stages between embryonic day 12 (E12) to E18 that required conducting time-mating. In those cases, we crossed either Calb2-CreERT2 or Ai14 females with males from the other strain and considered the vaginal plug day as E0. All mice were maintained on a 12h light cycle in the vivarium at Yale University. All protocols and procedures were approved by Yale University Animal Care and Use Committee.

### Injections of adeno-associated viruses (AAVs) into the transverse sinus

To label mitral/tufted cell bodies and axons across the cardinal axes of the OB, we injected recombinant adeno-associated viruses-9 (AAV9) driving the expression of the GCamP6 fluorescent calcium sensor, into the transverse sinus. This vector is commonly used to study neural activity that also triggers a robust expression of EGFP in projection neurons of the brain, including those from the OB (Barson et al., 2020; Hamodi et al., 2020). Injections were performed using a slightly modified version of the protocol described by Hamodi et al. (2020). Briefly, a solution of AAV9-syn-GCaMP6s (Addgene viral prep #[100843-AAV9](https://www.addgene.org/100843/)) at titer ≥ 1×10¹³ vg/mL was loaded in a Hamilton Neuros syringe (Hamilton #65460-06) attached to a 33-gauge beveled needle that was mounted in a stereotaxic frame. 6 CD1 pups at P0 (n = 6; 3 males, 3 females) were anesthetized by hypothermia on an ice-cold plate for 5 min. and then transferred to the stereotaxic frame on top of a cold metallic plate. The head of pups were fixed by the blunt portion of the ear bars, and the skin over the transverse sinuses was cut using microsurgical scissors (WPI #W16101). The needle was slowly advanced through the skull and the tip inserted into the transverse sinus, wherein 2 μL of the AAV9-syn-GCamP6 solution was injected bilaterally. The skin was then closed and sealed with 3M Vetbond Tissue Adhesive (1469SB). Following recovery on a heating pad, the pups were returned to the mother and maintained on a 12h light-dark cycle in the Yale vivarium until they were 33 days of age (P33).

### Injections of 4-Hydroxytamoxifen (4OHTx) on Calb2-CreERT2 × Ai14 mice

A 10 mg/mL solution of 4OHTx (Sigma-Aldrich #H6278) was prepared by dissolving 10 mg of 4OHTx in 100 μL of absolute ethanol on a rocker for 1 h at 55°C. Subsequently, 900 μL of sunflower oil (Sigma-Aldrich #S5007) was added for additional incubation of 30 min. at 37°C with constant shaking. Immediately afterward, the 4OHTx solution was loaded into insulin syringes, and each animal received two consecutive intraperitoneal (IP) injections of 40 mg/kg 4OHTx, separated 2 h apart, to achieve efficient Cre-mediated recombination. For adult tissues analyses at 10-14 weeks of age (n = 6, 3 males, 3 females), mice received a single injection on one postnatal day between P0 and P14. To obtain embryonic tissues at E12-E17 (n = 2 for E12; n = 3 for E13 to E17), pregnant females were injected at E11. No signs of distress or developmental defects were detected on injected pregnant females or their embryos that could have required humane end-point euthanasia.

### Tissue processing

Adult CD1 mice, CD1 mice injected with AAV at P33, and adult Calb2-CreERT2 × Ai14 mice were euthanized with an overdose of Euthasol (Virbac) and transcardially perfused with 5 mL of 0.9% saline followed by cooled 4% paraformaldehyde (PFA, EMS #19208) dissolved in phosphate-buffered saline (PBS) at a volume of 1 mL/g body weight. Brains were then extracted from the skulls and postfixed in PFA for 24 h. For embryonic tissues obtained from Calb2-CreERT2 × Ai14 mice, pregnant females at E12-E17 were euthanized in a CO2 chamber, and the embryos were rapidly extracted from the uterus. Six embryos from different dams were used per study point. Embryonic heads were collected, and an incision was made along the skull midline for subsequent fixation by immersion in PFA for 48 h at 4°C. After fixation, both adult and embryonic brain tissues were rinsed with PBS and cryoprotected in 30% sucrose-PBS. Once the tissues had sunk to the bottom of the sucrose solution, the brains or embryonic heads were embedded in Tissue-Tek OCT compound (Fisher Scientific #4585) and rapidly frozen for cryo-sectioning. Brains were serially sectioned with a Reichert Frigocut cryostat (E-2800) at 25 μm in the coronal plane and collected onto Fisherband ColorFrost Plus slides (Fisher Scientific). Sections were dried on a slide warmer at 50°C and stored at -80°C until further processing.

### Immunohistochemistry

In preparation for immunohistochemistry (IHC), sections were thawed on a slide warmer at 60°C for 20 min. and OCT was removed by washing in PBS for 10 min. Antigen retrieval was performed by immersing the slides in 0.01M citrate buffer (pH 6.0) at 65°C for 35 min., followed by a rapid transfer into the same ice-cooled buffer to induce a heat-shock. Slides were then placed horizontally into a humid chamber, and non-specific binding of antibodies was blocked by incubating the sections 1 h at room temperature (RT) in PBS containing 0.1% Triton X100 (AmericanBio #AB02025) (PBST), 5% Normal Donkey Serum (NDS, Southern Biotech #0030-01) or 5% Normal Goat Serum (NGS, Accurate Chemicals #CL1200-100), depending on the species of the secondary antibodies, and 0.1% Bovine Serum Albumin (BSA, Sigma Aldrich #A3059). Sections were incubated with primary antibodies (Table 1) diluted in a solution that is 1:10 dilution of the blocking solution for 24 h at 4°C. Sections were washed 3 times for 10 min. each with PBST and then incubated with the appropriate secondary antibodies (Table 1) diluted in PBST supplemented with 1 μg/mL DAPI (Sigma-Aldrich #D9542) for nuclear counterstaining. After washing 3 times for 10 min. each, sections were mounted with Mowiol 4-88 (Sigma-Aldrich #81381) before imaging.

**Table 1.** Antibodies.

| Antigen | Primary Ab | Source (Cat. #) | RRID | Dilution | Secondary Ab | Source | Dilution |
|---|---|---|---|---|---|---|---|
| Calretinin | Mouse IgG1 | Millipore-Sigma (MAB1568) | AB_94259 | 1:500 | Goat α-mouse IgG Alexa 555 | Thermo Fisher Sci. (A28180) | 1:500 |
|  |  |  |  |  | Donkey α-mouse IgG Alexa 488 | Jackson Immunores. (715-545-150) | 1:500 |
| Reelin (G10) | Mouse IgG1 | Abcam (ab78540) | AB_1603148 | 1:500 | Goat α-mouse IgG Alexa 488 | Thermo Fisher Sci. (A28175) | 1:1000 |
| Reelin | Rabbit clone [EPR26278-30] | Abcam (ab312310) | AB_3076463 | 1:500 | Goat α-rabbit IgG Alexa 488 | Thermo Fisher Sci. (A27034) | 1:2000 |
|  |  |  |  |  | Goat α-rabbit IgG Alexa 647 | Thermo Fisher Sci. (A21245) | 1:1000 |
| RFP | Rabbit Polyclonal | Rockland (600-401-379) | AB_2209751 | 1:500 | Donkey α-rabbit IgG Alexa 555 | Thermo Fisher Sci. (A31572) | 1:1000 |
| RFP | Goat Polyclonal | Rockland (200-401-379) | AB_2744552 | 1:1000 | Donkey α-goat IgG Alexa 555 | Thermo Fisher Sci. (A21432) | 1:1000 |
| Tbx21 | Rabbit Polyclonal | Gift from Yoshihiro Yoshihara (Yoshihara et al., 2005) |  | 1:1000 | Goat α-rabbit IgG Alexa 647 | Thermo Fisher Sci. (A27034) | 1:2000 |
|  |  |  |  |  | Donkey α-rabbit IgG Alexa 647 | BioLegend (406414) | 1:500 |
| Synaptophysin | Mouse IgM | BioLegend (837103) | AB_2783410 | 1:500 | Goat α-mouse IgM Alexa 488 | Thermo Fisher Sci. (A10680) | 1:1000 |

### Electron microscopy

For electron microscopy (EM), CD1 brain tissues were collected at embryonic stages E13 to E17 and postnatal ages P1, P9 and P21 (n = 3 per age). Embryos were obtained from pregnant females euthanized with CO_2_, rapidly removed from the uterus and their brains fixed for 4 days by immersion in a solution of ice-cold 4% PFA + 2% glutaraldehyde (GLA, EMS #16310) in PBS. For postnatal tissues, P1 mice were deeply anesthetized with hypothermia, whereas P9 and P21 mice were euthanized with Euthasol and transcardially perfused with 0.9% saline (1 mL for P9 mice and 5 mL for P21 mice), followed by fixative containing 4% PFA and 2% GLA (10 mL for P9 mice and 20 mL for P21 mice). Brains were then extracted from skulls and postfixed overnight on the same fixative solution. Subsequently, brains tissues were embedded in agarose (Lonza #50002) to obtain thick slices (50 µm) using a Vibratome 3000 system, prior to selecting PC regions of interest for micro-dissection. Tissues were post-fixed to preserve membrane lipids with 4% osmium (EMS #19160) for 1 h, dehydrated with graded alcohols, and stained with uranyl acetate (EMS #22400) and lead citrate 3% (EMS #22410). Subsequently, sections were embedded in EMbed 812 resin (EMS #14900) to obtain 40 nm ultra-thin sections using an UltraCut-E ultramicrotome from the anterior (APC) and posterior (PPC) piriform cortex. Microsections were mounted on formvar-coated slot grids (EMS #FF2010-CU) for imaging in a JEOL electron microscope (JEM-1200 EX II) at the level of the LOT in layer I of APC and PPC with 4800× and 2900× magnification respectively.

### Image acquisition and statistical analysis

All fluorescent labeling was imaged using a laser scanning confocal microscope (Zeiss LSM 800 with Airyscan) at different magnifications. To measure the extent of the low-CR-expressing region within the LOT (Fig. 1), two consecutive sections from the anterior PC were imaged at 10× magnification from three different CD1 mice (n = 3). Using ImageJ, the outlines of the LOT and the low-CR-expressing region were manually delineated with the Freehand Selection tool, and their respective areas were measured. The extent of the low-CR-expressing region was expressed as a percentage mean across the three animals of the total LOT area.

**Figure 1.**
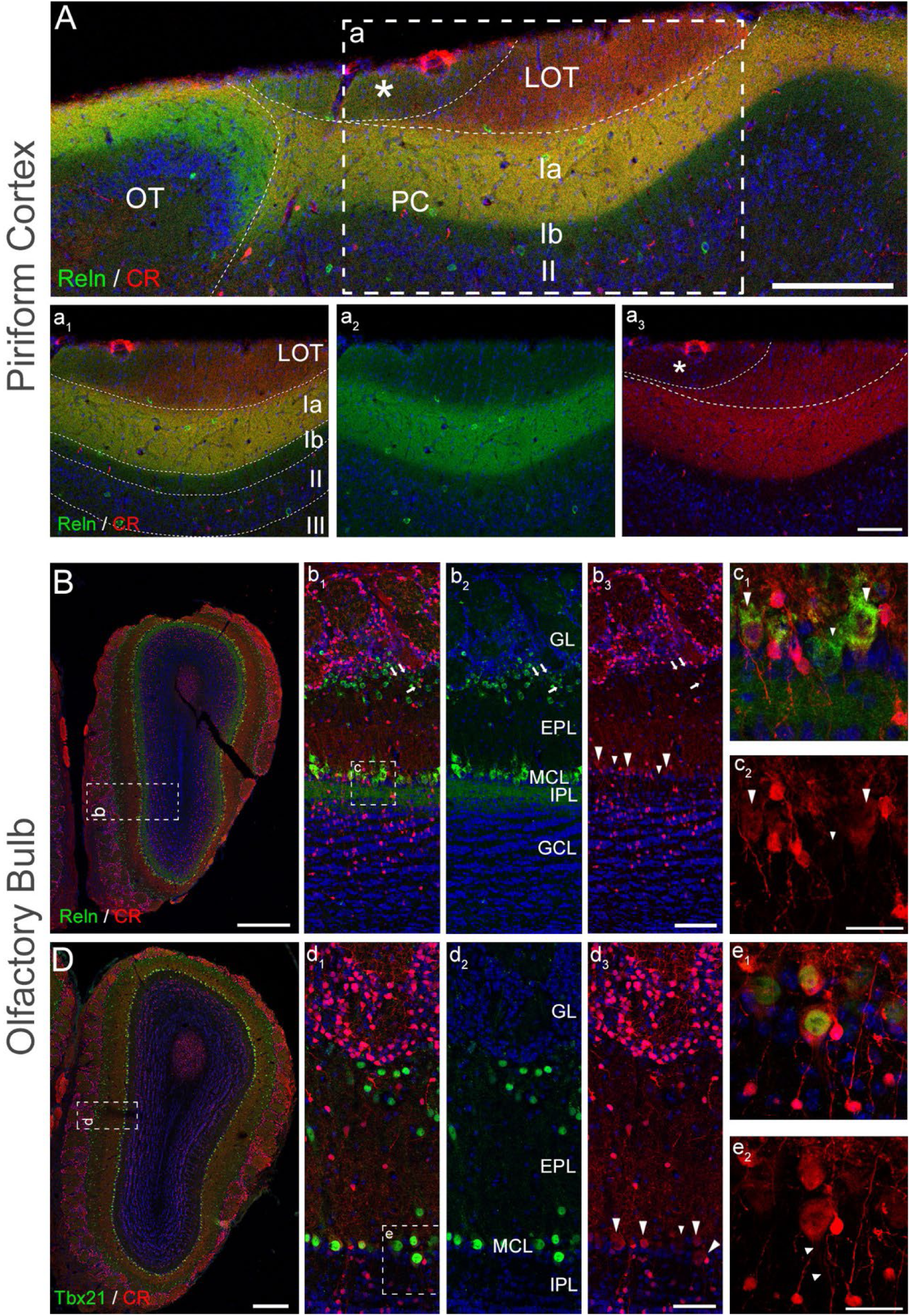
Immunohistochemistry to detect calretinin in both PC/OT tandem and OB of wild type CD1 mice. **(A, a)** Immunohistochemistry to colocalize CR (red, Mouse antibody) and Reln (green, Rabbit antibody) in PC and OT showing the layering with LOT axons expressing CR, layer Ia CR and Reln, and layers Ib and II no labeling for these molecules. A low-CR-labeling region next to the OT is seen in PC suggesting a topological organization of CR+ mitral/tufted cells in the OB. **(a1)** Intermediate magnification of PC layers, and **(a_2_–a_3_)** single-channel intermediate magnification of CR and Reln colocalization. **(B)** Low-magnification mosaic to colocalize CR (red) and Reln (green) in the OB showing labeling of mitral/tufted cells. **(b_1_)** Intermediate magnification of CR and Reln colocalization, and **(b_2_–b_3_)** single-channel plus DAPI intermediate magnification of CR and Reln colocalization. Mitral cells from the MCL show distinct patterns of CR expression with some having intense labeling (large arrowheads) while others have weak labeling (small arrowheads). Tufted cells are seen in the outer EPL (arrows). **(c_1_–c_2_)** High-magnification staining of CR and Reln colocalization in glutamatergic mitral cell neurons showing distinct levels of CR expression (large and small arrowheads). **(D)** Low-magnification mosaic to colocalize CR (red) and Tbx21 (green) in OB showing intense labeling of cells in the MCL and GL. **(d_1_)** Intermediate magnification of CR and Tbx21 colocalization labeling mitral/tufted cells. **(d_2_–d_3_)** single-channel plus DAPI intermediate magnification of CR and Tbx21 colocalization showing distinct levels of CR expression by mitral cells (large and small arrowheads). **(e_1_–e_2_)** High-magnification CR and Tbx21 colocalization in glutamatergic mitral cell neurons, some showing labeling of their axon hillock with CR (arrowheads). Notably, CR is also detected in a large subpopulation of interneurons in GCL and GL **(B, b_1_)**. Animals used were 8-10 weeks old. Abbreviations: CR, calretinin; EPL, external plexiform layer; GCL, granule cell layer; GL, glomerular layer; IPL, internal plexiform layer; LOT, lateral olfactory tract; MCL, mitral cell layer; OB, olfactory bulb; OT, olfactory tubercle; PC, piriform cortex; Reln, reelin. Scale bars: 500μm in **(B, D)**; 200μm in **(A)**; 100μm in **(a_3_, b_3_, d_3_)**; 50μm in **(c_2_, e_2_)**.

To determine the distribution of mitral/tufted cells that expressed CR and EGFP triggered by the AAVs injections along the cardinal axes of the OB, we quantified separately the number of mitral and tufted cells that were double labeled for CR^+^/Reln^+^, the last a marker for OB projection neurons (Alcantara et al., 1998; Ramos-Moreno et al., 2006), as well as EGFP^+^/Reln^+^, to assess for AAV infection efficiency. We imaged four regions of the OB from six different mice (n = 6; 3 males, 3 females): dorsal (d), ventral (v), medial (m) and lateral (l) at a 20× magnification (Fig. 2A), taken from three different sections of the middle region of the OB, separated 250 μm. Mitral cells were counted in the mitral cell layer (MCL), while tufted cells were counted within the outer plexiform layer (oEPL) in the periglomerular (PG) region. Since there is a subpopulation of PG neurons (PGNs) that express Reln but are not tufted cells (Okuyama-Yamamoto et al., 2005; Hellwig et al., 2012), we calculated the size of all PGN cell bodies within the oEPL to distinguish between PGN-tufted vs PGN-interneurons. The cell body areas were delineated using the ImageJ Freehand Selection tool and averaged between sections (replicates) and animals. Statistical differences were calculated using the GraphPad Prism 11 software and the applied tests are summarized in Table 2.

**Figure 2.**
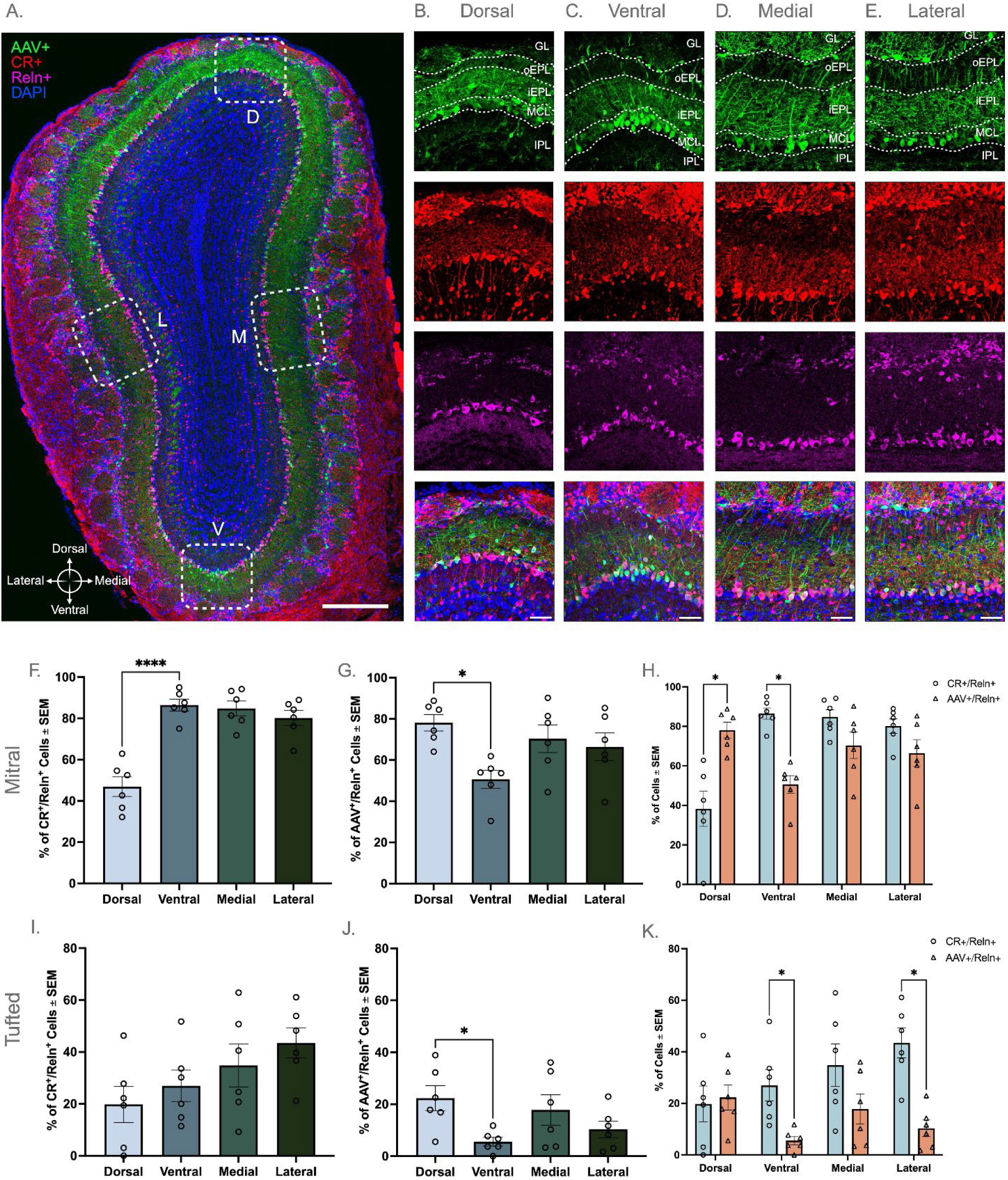
Distribution of CR-expressing mitral/tufted cells in OB along the cardinal axes. **(A)** Low-magnification mosaic of whole OB, AAV injected via transverse sinus in green, CR (Mouse antibody) in red, Reln (Rabbit antibody) to label OB projection neurons in magenta, and DAPI in blue. **(B)** High magnification of dorsal region, both single channels and merged. **(C)** High magnification of ventral region, both single channels and merged. **(D)** High magnification of medial region, both single channels and merged. **(E)** High magnification of lateral region, both single channels and merged. **(F–H)** Quantifications involving mitral cells: **(F)** Percentage of CR^+^/Reln^+^ cells across regions (one-way ANOVA with Tukey’s multiple comparisons) showing a significant lower percentage of double labeled neurons in the dorsal region, **(G)** Percentage of AAV^+^/Reln^+^ cells across regions (one-way ANOVA with Tukey’s multiple comparisons), showing a significant lower percentage of double labeled neurons in the ventral region, and **(H)** Comparison between CR and AAV efficacy across regions (multiple unpaired t-tests using Holm-Šídák method). **(I–K)** Quantifications involving tufted cells: **(I)** Percentage of CR^+^/Reln^+^ cells across regions (one-way ANOVA with Tukey’s multiple comparisons), **(J)** Percentage of AAV^+^/Reln^+^ cells across regions (one-way ANOVA with Tukey’s multiple comparisons), showing a significant lower percentage of neurons in the ventral region, and **(K)** Comparison between CR and AAV efficacy across regions (multiple unpaired t-tests using Holm-Šídák method). Animals used were 8-10 weeks old (n = 6; 3 males, 3 females). Abbreviations: AAV, adeno-associated viruses; CR, calretinin; D, dorsal; V, ventral; M, medial; L, lateral; GL, glomerular layer; iEPL, inner external plexiform layer; IPL, internal plexiform layer; oEPL, outer external plexiform layer; MCL, mitral cell layer; Reln, reelin. Statistical significance: *p < 0.05; ****p < 0.0001. Scale bars: 300μm in **(A)**; 50μm in **(B–E)**.

**Table 2.** Statistical results.

| Analyzed (Figure) | Test | Pairwise comparisons | Statistical value | Adjusted p Value |
|---|---|---|---|---|
| % CR<sup>+</sup>/Reln<sup>+</sup> in Mitral Cells (Fig. 2F) | One-way ANOVA + Tukey’s multiple comparison test | Dorsal vs Ventral | Mean Diff. -0.3956<br>95.00% CI of diff. -0.5457 to -0.2454 | <0.0001 |
|  |  | Dorsal vs Medial | Mean Diff. -0.3789<br>95.00% CI of diff. -0.5291 to -0.2288 | <0.0001 |
|  |  | Dorsal vs Lateral | Mean Diff. -0.3323<br>95.00% Cl of diff. -0.4824 to -0.1821 | <0.0001 |
| % AAV<sup>+</sup>/Reln<sup>+</sup> in Mitral Cells (Fig. 2G) | One-way ANOVA + Tukey’s multiple comparison test | Dorsal vs Ventral | Mean Diff. 0.2747<br>95.00% CI of diff. 0.05234 to 0.4971 | 0.0122 |
| % AAV<sup>+</sup>/Reln<sup>+</sup> in Tufted Cells (Fig. 2J) | One-way ANOVA + Tukey’s multiple comparison test | Dorsal vs Ventral | Mean Diff. 0.1673<br>95.00% CI of diff. 0.0008391 to 0.3338 | 0.0486 |
| % Mitral Cells Side-by-Side (Fig. 2H) | Multiple unpaired t-tests | Dorsal CR+/Reln+ vs AAV+/Reln+ | Difference. -0.3199<br>SE of Difference. 0.06223. df 10.00 | 0.001581 |
|  |  | Ventral CR+/Reln+ vs AAV+/Reln+ | Difference. 0.3583<br>SE of Difference. 0.05260. df 10.00 | 0.000187 |
| % Tufted Cells Side-by-Side (Fig. 2K) | Multiple unpaired t-tests | Ventral CR+/Reln+ vs AAV+/Reln+ | Difference. 0.2138<br>SE of Difference. 0.06296. df 10.00 | 0.020317 |
|  |  | Lateral CR+/Reln+ vs AAV+/Reln+ | Difference. 0.3314<br>SE of Difference. 0.06603. df 10.00 | 0.002087 |
| Tufted vs Periglomerular Soma Size (Suppl. Fig. 1) | Unpaired t-test | Tufted vs Periglomerular Soma Size | Mean Diff. -69.78<br>95.00% CI of diff. -81.83 to -57.74 | <0.0001 |
| LOT Axon Count (Fig. 5A) | One-way ANOVA + Tukey’s multiple comparison test | E13 vs E14 | Mean Diff. -1855899<br>95.00% CI of diff. -3318484 to -393314 | 0.0083 |
|  |  | E13 vs E15 | Mean Diff. -1774090<br>95.00% CI of diff. -3236675 to -311505 | 0.0121 |
|  |  | E13 vs E16 | Mean Diff. -1776927<br>95.00% CI of diff. -03239512 to -314342 | 0.0120 |
|  |  | E13 vs E17 | Mean Diff. -1720494<br>95.00% CI of diff. -3183079 to -257909 | 0.0155 |
|  |  | E13 vs P21 | Mean Diff. 1473519<br>95.00% CI of diff. 10934 to 2936104 | 0.0476 |
| Asymmetric Synapses Count (Fig. 6C) | Two-way ANOVA + Tukey’s multiple comparison test | P21 Anterior vs P21 Posterior | Mean Diff. -51320<br>95.00% CI of diff. -77974 to -24666 | <0.0001 |
|  |  | P21 Anterior vs P9 Anterior | Mean Diff. 46971<br>95.00% CI of diff. 20317 to 73625 | <0.0001 |
|  |  | P21 Anterior vs P9 Posterior | Mean Diff. 37125<br>95.00% CI of diff. 10471 to 63779 | 0.0011 |
|  |  | P21 Anterior vs P1 Anterior | Mean Diff. 56375<br>95.00% CI of diff. 29721 to 83029 | <0.0001 |
|  |  | P21 Anterior vs P1 Posterior | Mean Diff. 67547<br>95.00% CI of diff. 40893 to 94201 | <0.0001 |
|  |  | P21 Anterior vs E17 Anterior | Mean Diff. 60179<br>95.00% CI of diff. 33525 to 83029 | <0.0001 |
|  |  | P21 Anterior vs E17 Posterior | Mean Diff. 68241<br>95.00% CI of diff. 41587 to 94895 | <0.0001 |
|  |  | P21 Anterior vs E16 Anterior | Mean Diff. 65822<br>95.00% CI of diff. 39168 to 92476 | <0.0001 |
|  |  | P21 Anterior vs E16 Posterior | Mean Diff. 66622<br>95.00% CI of diff. 39968 to 93276 | <0.0001 |
|  |  | P21 Anterior vs E15 Anterior | Mean Diff. 57877<br>95.00% CI of diff. 31223 to 84531 | <0.0001 |
|  |  | P21 Anterior vs E15 Posterior | Mean Diff. 70265<br>95.00% CI of diff. 43611 to 96919 | <0.0001 |
|  |  | P21 Anterior vs E14 Anterior | Mean Diff. 55947<br>95.00% CI of diff. 29293 to 82601 | <0.0001 |
|  |  | P21 Anterior vs E14 Posterior | Mean Diff. 68430<br>95.00% CI of diff. 41776 to 95084 | <0.0001 |
|  |  | P21 Anterior vs E13 Anterior | Mean Diff. 74033<br>95.00% CI of diff. 47379 to 100687 | <0.0001 |
|  |  | P21 Anterior vs E13 Posterior | Mean Diff. 74736<br>95.00% CI of diff. 48082 to 101390 | <0.0001 |
|  |  | P21 Posterior vs P9 Anterior | Mean Diff. 98291<br>95.00% CI of diff. 71637 to 124945 | <0.0001 |
|  |  | P21 Posterior vs P9 Posterior | Mean Diff. 88445<br>95.00% CI of diff. 61791 to 115099 | <0.0001 |
|  |  | P21 Posterior vs P1 Anterior | Mean Diff. 107695<br>95.00% CI of diff. 81041 to 134349 | <0.0001 |
|  |  | P21 Posterior vs P1 Posterior | Mean Diff. 118867<br>95.00% CI of diff. 92213 to 145521 | <0.0001 |
|  |  | P21 Posterior vs E17 Anterior | Mean Diff. 111499<br>95.00% CI of diff. 84845 to 138153 | <0.0001 |
|  |  | P21 Posterior vs E17 Posterior | Mean Diff. 119561<br>95.00% CI of diff. 92907 to 146215 | <0.0001 |
|  |  | P21 Posterior vs E16 Anterior | Mean Diff. 117142<br>95.00% CI of diff. 90488 to 143796 | <0.0001 |
|  |  | P21 Posterior vs E16 Posterior | Mean Diff. 117942<br>95.00% CI of diff. 91288 to 144596 | <0.0001 |
|  |  | P21 Posterior vs E15 Anterior | Mean Diff. 109197<br>95.00% CI of diff. 82543 to 135851 | <0.0001 |
|  |  | P21 Posterior vs E15 Posterior | Mean Diff. 121585<br>95.00% CI of diff. 94931 to 138239 | <0.0001 |
|  |  | P21 Posterior vs E14 Anterior | Mean Diff. 107267<br>95.00% CI of diff. 80613 to 133921 | <0.0001 |
|  |  | P21 Posterior vs E14 Posterior | Mean Diff. 119750<br>95.00% CI of diff. 93096 to 146404 | <0.0001 |
|  |  | P21 Posterior vs E13 Anterior | Mean Diff. 125353<br>95.00% CI of diff. 98699 to 152007 | <0.0001 |
|  |  | P21 Posterior vs E13 Posterior | Mean Diff. 126056<br>95.00% CI of diff. 99402 to 152710 | <0.0001 |
|  |  | P9 Anterior vs E13 Anterior | Mean Diff. 27062<br>95.00% CI of diff. 408.3 to 53716 | 0.0437 |
|  |  | P9 Anterior vs E13 Posterior | Mean Diff. 27765<br>95.00% CI of diff. 1111 to 54419 | 0.0345 |
|  |  | P9 Posterior vs P1 Posterior | Mean Diff. 30422<br>95.00% CI of diff. 3768 to 57076 | 0.0137 |
|  |  | P9 Posterior vs E17 Posterior | Mean Diff. 31116<br>95.00% CI of diff. 4462 to 57770 | 0.0106 |
|  |  | P9 Posterior vs E16 Anterior | Mean Diff. 28697<br>95.00% CI of diff. 2043 to 55351 | 0.0251 |
|  |  | P9 Posterior vs E16 Posterior | Mean Diff. 29497<br>95.00% CI of diff. 2843 to 56151 | 0.0190 |
|  |  | P9 Posterior vs E15 Posterior | Mean Diff. 33140<br>95.00% CI of diff. 6486 to 59794 | 0.0050 |
|  |  | P9 Posterior vs E14 Posterior | Mean Diff. 31305<br>95.00% CI of diff. 4651 to 57959 | 0.0099 |
|  |  | P9 Posterior vs E13 Anterior | Mean Diff. 36909<br>95.00% CI of diff. 10255 to 63563 | 0.0012 |
|  |  | P9 Posterior vs E13 Posterior | Mean Diff. 37611<br>95.00% CI of diff. 10957 to 64265 | 0.0009 |
| Symmetric Synapses Count (Fig. 6D) | Two-way ANOVA + Tukey’s multiple comparison test | P21 Posterior vs E17 Anterior | Mean Diff. -16581<br>95.00% CI of diff. -32069 to -1092 | 0.0266 |

For the EM analysis, we measured two different parameters across the embryonic development from E13 to E17 and postnatally at P1, P9 and P21: 1) the number of LOT axons in the anterior PC (APC), and 2) the density of asymmetric (excitatory) and symmetric (inhibitory) synapses in layer Ia of both APC and PPC. To quantify the number of LOT axons, we took 8 randomly selected images at 4800× from the LOT region from three different animals per study point. Numbers were averaged between replicates and animals and differences calculated using GraphPad Prism 11 (Table 2 for statistical analysis). To calculate the density of synapses, we used 8 randomly selected images from layer Ia of both APC and PPC at 2900× taken from three different animals per study point. Asymmetric synapses (excitatory) were recognized by their larger size, the presence of thick electrodense accumulation at the postsynaptic terminal and round vesicles in the presynaptic terminal, while symmetric synapses (inhibitory) were identified by their smaller size, thin pre- and postsynaptic densities that looked relatively similar on both sides of the membrane, and often oval/flattened presynaptic vesicles. Comparisons between both types of synapses in APC and PPC, as well as across ages, were measured using GraphPad Prism 11 and used statistical tests and significant differences are represented in Table 2:

## Results

### Calretinin labels mitral and tufted cells in the olfactory bulb

CR is a calcium-binding protein expressed by 10-30% of GABAergic interneurons across the brain (Barinka and Druga, 2010; Cauli et al., 2014). In the olfactory system, CR is expressed in interneurons, where it regulates centrifugal input from the basal forebrain (Constantinescu et al., 2025). Of interest, CR is also expressed by the glutamatergic OB projection neurons, mitral/tufted cells (Jacobowitz and Winsky, 1991; Resibois and Rogers, 1992; Wouterlood and Härtig, 1995; Pombal et al., 2002). Despite these observations, the expression of CR by the OB projection neurons has received little to no attention over the years other than to visualize the LOT and distinguishing layers in PC (Sarma et al., 2011; Amaniti et al., 2015; Martin-Lopez et al., 2019a; Martin-Lopez et al., 2019b). In the present study, we first characterized the expression of CR throughout the olfactory system, with particular emphasis on projection neurons.

To ensure that CR labeling corresponded to projection neurons, we performed double immunolabeling for CR and Reelin (Reln), an established marker for mitral/tufted cells and their axons in the LOT (Alcantara et al., 1998; Martinez-Cerdeno et al., 2003; Ramos-Moreno et al., 2006; Martin-Lopez et al., 2011). The double labeling revealed three main findings: First, both CR and Reln labeled the LOT but with poor resolution of individual axons (Fig. 1A, a_1_-a_3_) (Martin-Lopez et al., 2019a; Martin-Lopez et al., 2019b). Second, the labeling for both CR and Reln was more intense in layer Ia of PC, where LOT axons make synapses with apical dendrites of PC pyramidal cells, suggesting a higher concentration of both CR and Reln in axon terminals. Third, we identified a region of low-CR-labeling ventrally at the boundary of the LOT, proximal to the OT (Fig. 1A, a_3_, asterisk). This region represented approximately 37% of the total LOT area and was not previously recognized but suggested a differential expression of CR by mitral/tufted cells that could reflect topographic specificity. Using both Reln and Tbx21 as previously documented co-markers of mitral/tufted cells (Yoshihara et al., 2005; Imamura et al., 2011) we found that CR was expressed in mitral cells in the OB mitral cell layer (MCL) (Fig. B-c_1_, D-e_1_), across the circumference of the OB with different intensities (Fig. 1B, b_1_-b_3_, c_2_, d_3_, large vs small arrowheads). Interestingly, the CR immunostaining did not reveal individual dendrites and/or axons in mitral cells, except occasionally in the region of the axon hillock (Fig. 1e_2_, arrowheads), highlighting different CR content among OB neurons. Similarly, in the outer external plexiform layer (oEPL) CR labeled tufted cells were intermingled with several interneuron subpopulations interneurons in the juxtaglomerular region (Fig. 1b_1_-b_3_, d_1_-d_3_, arrows).

Building on these observations and previous work suggesting molecular specificity of projection neurons in OB domains (Treloar et al., 2002; Walz et al., 2006; Imamura et al., 2011; Imamura and Greer, 2015), we next asked whether CR expression in mitral/tufted cells exhibited a topographic distribution along the cardinal axes of the OB that could account for the differential CR labeling observed in the LOT. Because CR immunohistochemistry (IHC) did not reveal axons, we sought to compare CR expression with a reporter expressed by OB projection neurons to assess distribution and CR expression of individual axons within the LOT. To this end, we performed IHC for CR and Reln on OB sections from mice that were previously injected into the transverse sinuses with AAVs (Hamodi et al., 2020), a procedure used to drive EGFP expression in OB projection neurons and track their axons into the LOT. We quantified both mitral and tufted cells separately expressing either CR or AAV-EGFP in four different regions of the OB—dorsal, medial, ventral and lateral (Fig. 2A) in combination with Reln as a proxy to differentiate projection neurons from interneurons. Quantification of cells in the MCL revealed that more than 80% of mitral cells expressed CR, identified as CR^+^/Reln^+^ neurons, across the ventral (86%), medial (85%) and lateral (80%) regions (Fig. 2F). However, this percentage significantly decreased in the dorsal area of the OB, where only 47% of neurons were CR^+^/Reln^+^ (Fig. 2B-E, F), suggesting that the low-CR-labeling region of the LOT may be accounted for in part by the lack of CR^+^ axons from mitral cells from the dorsal OB. Furthermore, the percentage of mitral cells that were efficiently infected with AAVs (AAV^+^/Reln^+^) showed that viral infections of mitral cells were heterogenous across the circumference of the OB and in all regions lower than 80% of the total number of mitral cells. Thus, the dorsal region showed a 78% infection rate efficiency, 70% for the medial, 66% for the lateral and a decline to 51% for the ventral regions, respectively (Fig. B-E, G). While the AAV injections did effectively label the soma, dendrites and axons of infected mitral cells, the AAV injections may not fully label all OB mitral populations (Fig. 2H).

For tufted cells analysis, we quantified only those Reln^+^ neurons located in the outer external plexiform layer (oEPL), adjacent to the OB glomeruli, that met a predefined soma-size criterion. This criterion was necessary because the oEPL contains both tufted cells and periglomerular neurons (PGNs), some of which also express Reln (Okuyama-Yamamoto et al., 2005; Hellwig et al., 2012). To establish a size threshold, we performed IHC for Reln and Tbx21 in a separate set of tissue sections and measured the soma area of Reln^+^ neurons classified as tufted cells (Reln^+^/Tbx21^+^) or PGN (Reln^+^/Tbx21^-^) neurons. Tufted cells exhibited a significantly larger mean soma area (142.00μm^2^) relative to PGNs (72.24μm^2^) (Suppl. Fig. 1; end of this document). Using a soma area cutoff of 100.00μm^2^ for inclusion, we established a reliable criterion for identifying tufted cells in the oEPL. We then quantified the proportion of CR^+^/Reln^+^ tufted cells across the cardinal axes of the OB. Compared with mitral cells, a substantially smaller proportion of tufted cells expressed CR; 20% in dorsal, 27% in ventral and 35% in the medial, and 43% in the lateral regions (Fig. 2B-E, I). We next assessed AAV labeling efficiency by quantifying the proportion of Reln^+^ tufted cells that were also AAV^+^. The percentages were 22% for dorsal, 5% for ventral, 18% for medial and 10% in the lateral regions, with significantly lower labeling efficiency in the ventral region (Fig. 2J). Notably, AAV^+^/Reln^+^ tufted cells were less abundant than CR^+^/Reln^+^ tufted cells in the ventral and lateral domains (Fig. 2K). Consistent with the analysis of mitral cells, it appears that the AAV injections were not effective for labeling the entire population of tufted cells. There were no differences in percentages of mitral/tufted cells from males and females; therefore, data from both sexes were pooled for all subsequent analyses (Suppl. Fig. 2; end of this document). Together, these results showed a markedly lower CR labeling for mitral/tufted cells in the dorsal region of the OB, likely accounting for the reduced CR labeling observed in the ventral LOT in APC, which we termed the low-CR-labeling region. Additionally, AAV-mediated labeling following transverse sinus injection was variable across OB projection neurons, limiting the utility of this approach for comprehensively quantifying the developmental profile of the mitral/tufted cell population across the different OB domains.

### Tracking mitral and tufted cells axons with Calb2-CreERT2 × Ai14

Although CR is expressed by mitral/tufted cells in the OB IHC does not provide resolution of axonal processes. Thus, we searched for an alternative tool that would ensure labeling of CR^+^ neurons in their entirety enabling us to trace their axonal projections during LOT development. We evaluated Calb2-CreERT2 × Ai14 mice as a model for labeling neuronal processes, taking advantage of the robust expression of the fluorophore tdTomato in CR^+^ neurons after injections of 4OHTx (Fig. 3A). To validate this model for studying OB connectivity, we examined adult mice (10-14 weeks old) that had received injections of 4OHTx between P0 to P14. Our analysis showed a robust labeling of mitral/tufted cells and their individual axons within the LOT, which extended into both PC and OT (Fig. 3B, b_1_-b_3_). Labeled axons were also evident in layer Ia of PC, where LOT axons form synapses with apical dendrites of layers II/III pyramidal neurons (Bekkers and Suzuki, 2013). As expected, tdTomato^+^ axons colocalized with the protein CR detected by IHC (Fig. 3b_1_). However, unlike CR IHC, tdTomato labeling was relatively uniform throughout the LOT, with no apparent low-CR-labeling region in its ventral portion (Fig. 3B, b_1_–b_3_). This discrepancy likely reflects the different readouts of the two labeling approaches: CR IHC reveals the distribution of CR protein and is therefore sensitive to regional differences in its abundance, whereas tdTomato expression is driven by CreERT2-mediated recombination of the Ai14 reporter allele. Once recombination occurs, tdTomato expression is largely independent of the endogenous CR protein level. Thus, even if CR expression varies among mitral/tufted cells projecting to different LOT regions, these differences would not be reflected in tdTomato labeling. We, therefore, interpreted the low-CR-labeling region detected by IHC as potentially reflecting lower CR protein abundance in axons within the ventral LOT, rather than a reduced density of axons themselves.

**Figure 3.**
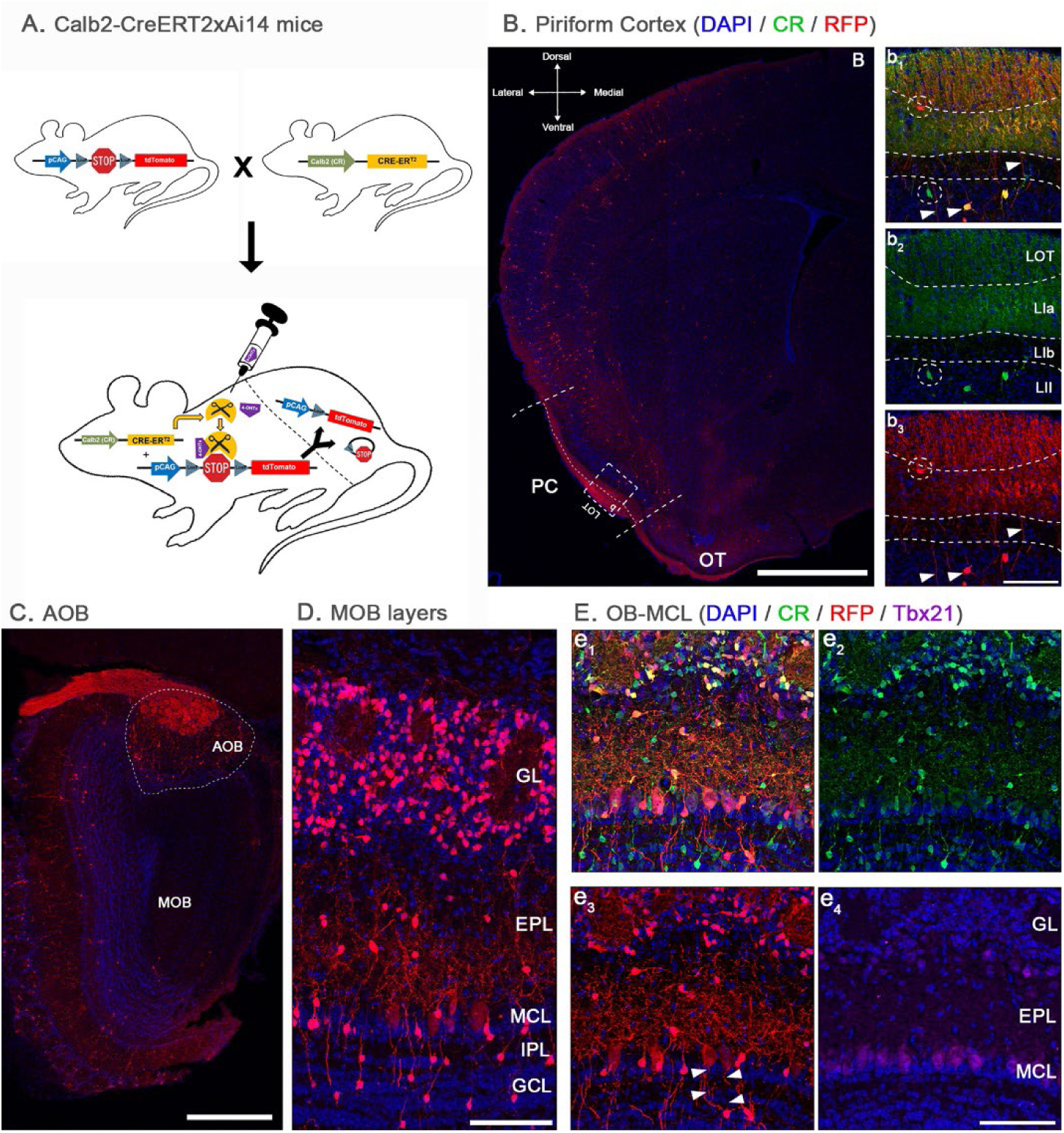
Mouse model to track neurons and axons of CR-expressing neurons in embryonic development. **(A)** Calb2-CreERT2 × Ai14 mouse genotype diagram. **(B)** Low-magnification mosaic to colocalize CR (green, Mouse antibody) and RFP (red, Rabbit antibody; reporter protein after injection with 4OHTx) in PC showing the LOT axons. **(b_1_)** Intermediate magnification of CR and RFP colocalization, and **(b_2_–b_3_)** single-channel plus DAPI intermediate magnification of CR and RFP colocalization. LOT axons are visible (arrowheads) and occasionally some interneurons were immunolabeled with the CR antibody but not the fluorophore and vice versa (dotted circles). **(C)** Low-magnification mosaic to colocalize the expression of the tdTomato fluorophore and immunolabeling with RFP in the AOB (dotted line). **(D)** Intermediate magnification showing the expression of the fluorophore tdTomato and immunolabeling with RFP in the MOB showing multiple periglomerular neurons and granule cells, as well as mitral/tufted cells and their processes. **(E, e_1-4_)** High magnification of the OB to colocalize CR (green, Mouse antibody), RFP (red, Goat antibody), and Tbx21 (magenta, Rabbit antibody) in the MCL, confirming expression in mitral/tufted cells projection neurons. Mitral cells show clear labeling of their processes and axons (**e_3_**, **arrowheads**). Animals used were 10-14 weeks old. Abbreviations: AOB, accessory olfactory bulb; CR, calretinin; EPL, external plexiform layer; GCL, granule cell layer; GL, glomerular layer; IPL, internal plexiform layer; LOT, lateral olfactory tract; MCL, mitral cell layer; MOB, main olfactory bulb; OB, olfactory bulb; OT, olfactory tubercle; PC, piriform cortex; Reln, reelin. Scale bars: 1mm in **(B)**; 500μm in **(C)**; 200μm in **(D)**; 100μm in **(b_3_, e_4_)**.

In the OB, tdTomato labeling was strong in the olfactory nerve (ON) axons innervating the glomeruli of the accessory olfactory bulb (AOB), whereas comparable labeling was not observed in the main olfactory bulb (MOB). This pattern suggested that CR may be preferentially expressed by OSNs originating in the vomeronasal organ (VNO), although further investigation of this system falls outside the scope of the present study. Within the MOB, tdTomato labeled mitral/tufted cells in the MCL and oEPL, consistent with the CR IHC results (Fig. 3D). Additionally, distinct subpopulations of PGNs in the glomerular layer (GL) and superficial granule cells in the granule cell layer (GCL) exhibited strong tdTomato expression (Fig. 3D), consistent with the well-characterized CR^+^ interneuron populations of the adult OB (Capsoni et al., 2021). The identity of mitral cells in the MCL was confirmed by their co-expression of tdTomato and Tbx21 (Fig. 3D, magenta), which was not present in interneurons. Consistent with the pattern observed by CR IHC, tdTomato labeling in mitral/tufted cells was weaker than that in interneurons (Fig. 3D, green). Calb2-CreERT2 × Ai14 mice also displayed a robust labeling of mitral/tufted cell axons in the OB, which could be readily traced through the LOT (Fig. 3e_3_, arrowheads). Together, these findings demonstrated that the Calb2-CreERT2 × Ai14 mice are well suited for studying the morphology of mitral/tufted cells and tracing their connectivity in PC through the LOT.

### Development of the lateral olfactory tract (LOT)

We next investigated the development of the LOT in Calb2-CreERT2 × Ai14 embryos, beginning at embryonic day 12 (E12), the earliest age at which mitral cell differentiation is detected in the OB (Lopez-Mascaraque and de Castro, 2002; Blanchart et al., 2006; Imamura et al., 2011). We tracked the mitral/tufted cells axons along the anterior-posterior axis to include all central regions of the olfactory pathway that we divided as follows: 1) the OB, to study the differentiation of mitral/tufted cells in the primordium of the OB; 2) the anterior olfactory nucleus (AON), which represents the first cortical relay station of the olfactory pathway (Martin-Lopez et al., 2025); and 3) piriform cortex (PC), which we subdivided from anterior to posterior regions as: a) anterior PC (APC), where layer II neurons curve to accommodate the LOT; b) intermediate PC (IPC), corresponding to the area where interhemispheric axons from the anterior commissure cross to the contralateral side; and c) posterior PC (PPC), corresponding to the associative cortex located next to the hippocampus where the LOT is almost invisible making PC layers II/III form a straight line (Chen et al., 2014; Martin-Lopez et al., 2019a). Next to PPC is the AMY, part of the limbic system that was also studied while analyzing this area.

At E12, we found pioneer mitral cells with radial orientation in the primordium of the OB (Fig. 4A_1_, MC insets) (Hinds, 1972; Hinds and Ruffett, 1973; Blanchart et al., 2006; Imamura et al., 2011). Beyond the OB we found evidence of tdTomato labeling throughout the entire length of the ventrolateral area from the lateral ganglionic eminence (LGE) (Martin-Lopez et al., 2019a; Martin-Lopez et al., 2025), which we interpreted as presumptive mitral cell axons that reached the anterior-posterior extent of the developing LOT region (Fig. 4A_2-5_, arrowheads). The presence of tdTomato labeling in posterior regions of the brain, proximal to the third ventricle and the caudal ganglionic eminence (CGE) suggested the formation of the LOT occurred earlier than previously suggested (Hinds, 1972; Sato et al., 1998; Lopez-Mascaraque and de Castro, 2002; Walz et al., 2006). No synaptophysin labeling, which we used as a proxy for synapse formation, was seen at E12. Our data showed that CR expression begins remarkably early in a subset of mitral cells, coinciding with the onset of their differentiation.

**Figure 4.**
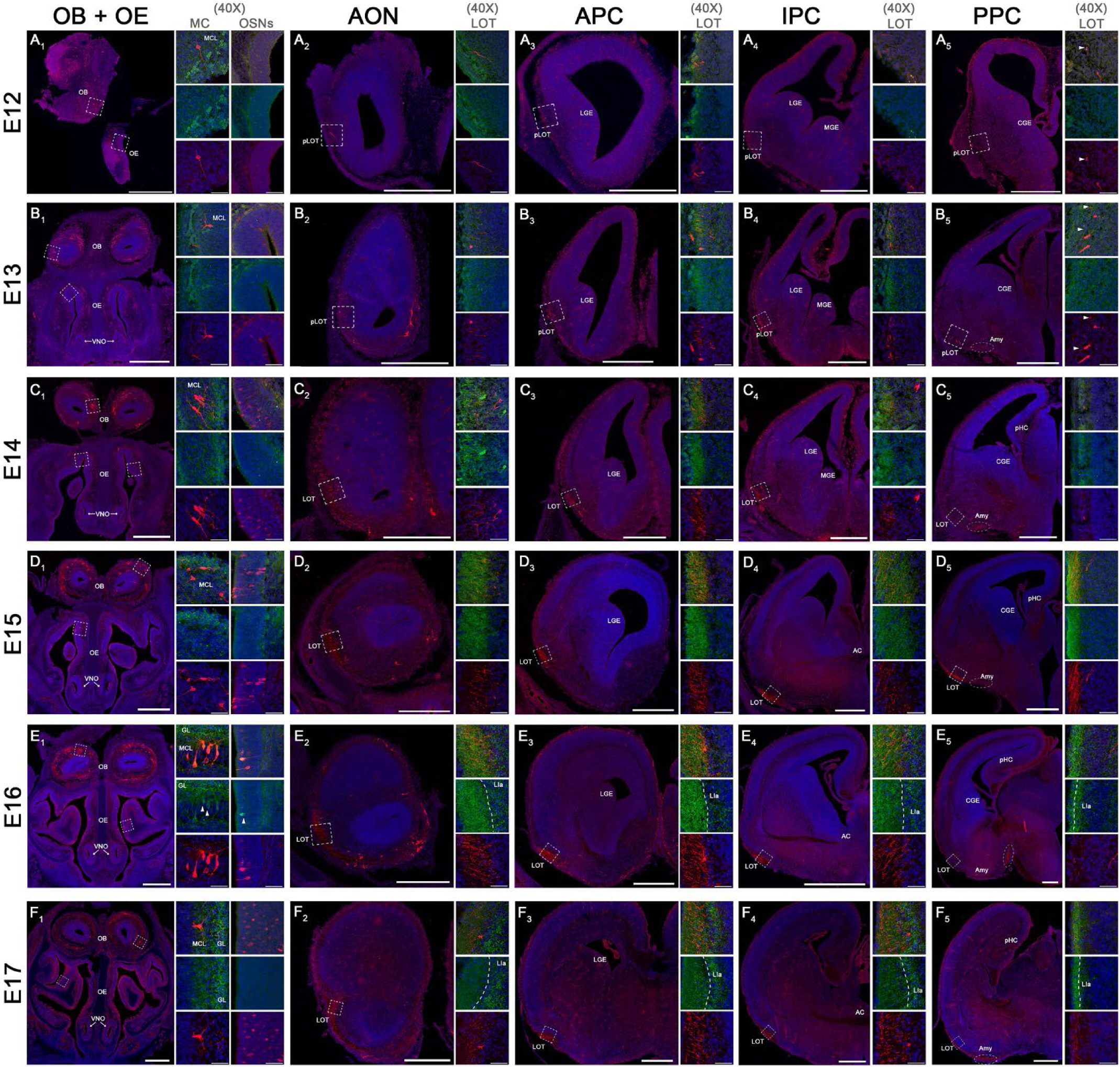
Embryonic development of LOT axons using CR as proxy. In all figures, synaptophysin (Mouse antibody) is green, RFP (Rabbit antibody; reporter protein after injection with 4OHTx) is in red, and DAPI in blue. **(A-F)** Sequence of high- and low-magnification images of the olfactory system in the OB/OE, AON, APC, IPC, and PPC across the embryonic development from E12 to E17. **(A_1–5_)** E12 developmental stage showing pioneer mitral cells and growth cones from the LOT reaching the PPC. No labeling for synaptophysin is visible. **(B_1-5_)** E13 stage showing dendritic arbors in some mitral cells and first OSNs labeled with RFP. There is a thickening of the LOT reaching the amygdala in posterior areas (dotted circle) and some growth cones arriving at the PPC (arrowheads). **(C_1–5_)** E14 stage showing a thicken LOT from the AON to the IPC, and only growth cones reaching the PPC. **(D_1–5_)** E15 developmental stage is the first one at which the LOT is visible in layer I of PPC and amygdala with expression of synaptophysin. Strong labeling of OSNs in the dorsal OE is evident. **(E_1–5_)** E16 shows strong labeling of layer Ia of PC with synaptophysin suggesting synaptic maturation. Synaptophysin is found labeling mitral cells and OSNs, and this age is the first evidence of VNO sensory neurons labeling with RFP-CR. **(F_1–5_)** E17 shows full maturation of the LOT across the entire anterior-posterior axis with strong labeling for synaptophysin. Abbreviations: Amy, amygdala; AC, anterior commissure; AON, anterior olfactory nucleus; APC, anterior piriform cortex; CGE, caudal ganglionic eminence; GL, glomerular layer; IPC, intermediate piriform cortex; LGE, lateral ganglionic eminence; LOT, lateral olfactory tract; MC, mitral cell; MCL, mitral cell layer; MGE, medial ganglionic eminence; OB, olfactory bulb; OE, olfactory epithelium; OSN, olfactory sensory neuron; pLOT, prospective lateral olfactory tract; pHC, prospective hippocampus; PPC, posterior piriform cortex. Scale bars: 500μm in all low-magnification images; 50μm in all accompanying high-magnification images.

**Figure 5.**
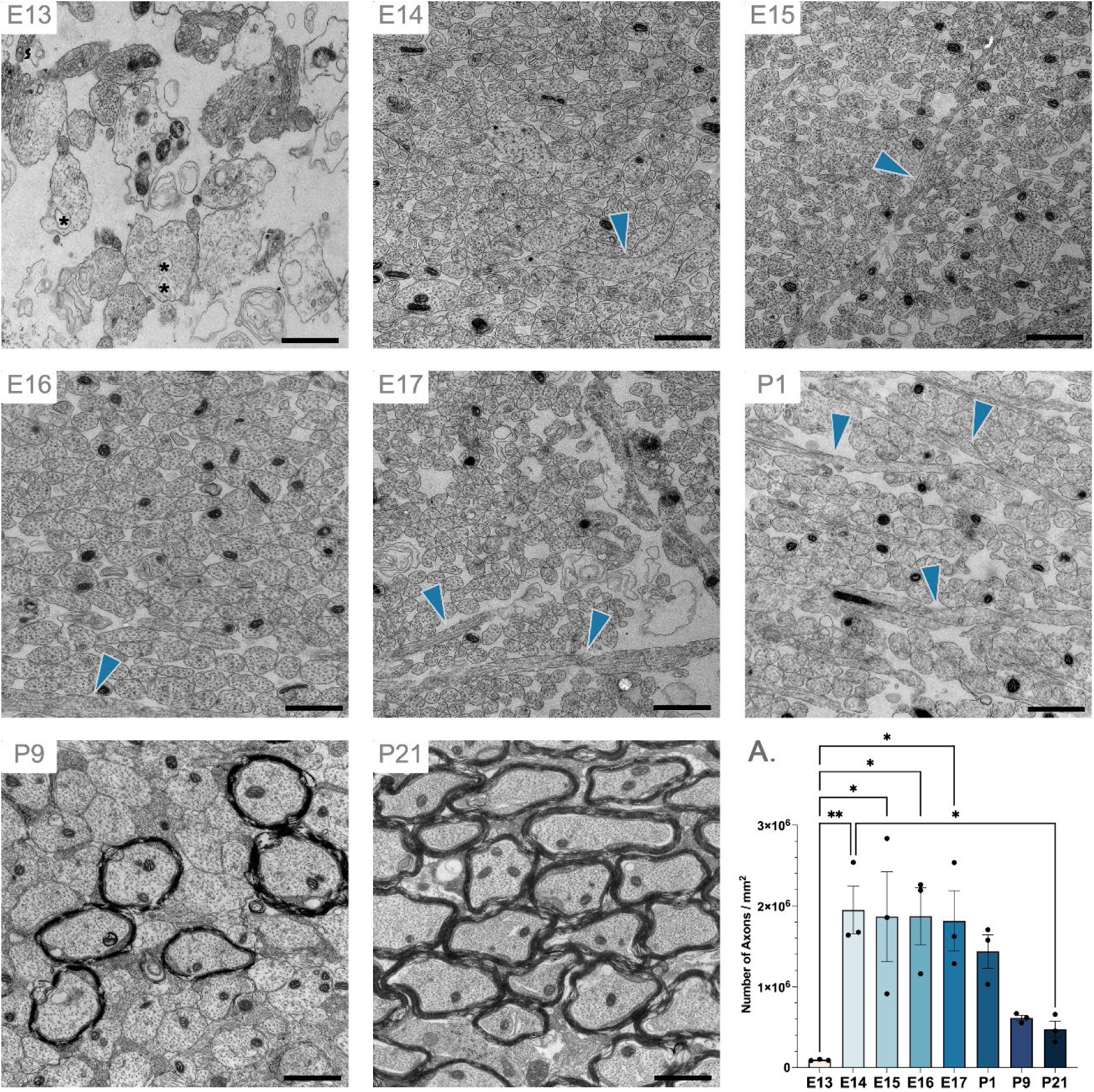
EM shows ultrastructural maturation of LOT axons across embryonic and early postnatal development. Representative EM images selected from each age E13–E17, P1, P9, and P21. Blue arrowheads represent axon collaterals. **(A)** Quantification of number of LOT axons per mm^2^ throughout development. One-way ANOVA with Tukey’s multiple comparisons. Statistical significance: *p < 0.05; **p < 0.01. Scale bars: All 1μm across ages.

By E13, the primordium of the OB showed differentiated mitral cells extending arborized apical dendrites to the region of the proto-glomeruli (Fig. 4B_1_, MC insets). By this age more tdTomato-labeled axons arrived at the AON, APC and IPC regions which collectively increased the width of the LOT (Fig. 4B_2-4_). Near posterior brain regions, most LOT axons were seen reaching the presumptive AMY (Fig. 4B_5_, dotted circle) with a few reaching the PPC. We first detected the synaptophysin expression (green labeling, Fig. 4B_2-5_), indicative of synaptogenesis, at E13.

From E14 onwards, the development of the LOT progressed more steadily as mitral cells continued to differentiate and changed their orientation from tangential-to-radial (Fig. 4C_1_, MC insets), a typical signature of maturation (Hinds and Ruffett, 1973; Blanchart et al., 2006). In addition, tufted cell neurogenesis is robust at E14 and is contributing axons to the LOT (Hinds, 1968a; Imamura et al., 2011; Hirata et al., 2019). The LOT continued to increase in thickness as more axons extended towards the IPC and elongation of axons continued (Fig. 4C_2-4_), showing initial collateralization into PC (Hirata and Fujisawa, 1999). As noted above, in the PPC most axons seemed to end in the AMY with fewer fibers reaching the PPC (Fig. 4C_5_). Immunolabeling for synaptophysin increased throughout the PC beginning at E14 suggesting the onset of functional maturation.

At E15 the differentiation of mitral cells in the OB was evident with apical dendrites forming a proto-glomerular layer (Fig. 4D_1_, MC insets). Concurrently, the LOT continued to increase in width as more axons joined the tract (Fig. 4D_2-4_). At this stage the axon innervation was prominent in PPC alongside the ongoing innervation of the AMY (Fig. 4D_5_). These observations suggested that the AMY is more precocious in development and precedes the innervation of PPC by mitral/tufted cell axons. By E16, although mitral/tufted cell neurogenesis is nearly complete (Imamura et al., 2020), ongoing maturation of OB projection neurons was still evident (Fig. 4E_1_, MC insets). The LOT was now clearly established along the entire rostrocaudal extent of PC, with evident axonal collateralization and branching within layer Ia (Fig. 4E_2-5_). This pattern was accompanied by strong synaptophysin labeling in layer Ia, consistent with the progressive establishment of presynaptic terminals in the PC (Fig. 4E_2-5_).

At E17 the OB showed characteristics of mitral/tufted cell maturation with a strong labeling (green) of synaptophysin in the developing glomerular layer (Fig. 4F1, MC insets). Compared to adults (Fig. 3 for reference), the number of tdTomato^+^ cells in the MCL was significantly lower, indicating that the expression of CR by many mitral/tufted cells likely occurred gradually and continued postnatally. In the LOT, axon collaterals extended into layer Ia of PC in all regions along the anterior-posterior axis, all showing an intense, dotted labeling with synaptophysin in layer Ia of PC (Fig. 4F_2-5_). Collectively, these data provide a complete map and comprehensive analysis of the LOT development and mitral/tufted cells differentiation/maturation, using the expression of CR as a proxy to track axonal growth from the OB. We established the onset of axogenesis for CR^+^ mitral cells at E12 and studied the sequential innervation of PC and AMY. Because the EPL in the prospective OB is difficult to distinguish from the MCL at embryonic stages (Blanchart et al., 2006), we could not definitively determine the contribution of tufted cells to these axonal projections. However, given that tufted cells do not innervate the PPC (Nagayama et al., 2010; Igarashi et al., 2012; Chon et al., 2020), we reasoned that the axons reaching the PPC and AMY in our analysis were predominantly, if not exclusively, derived from mitral cells.

Although the peripheral olfactory system was outside the scope of our analysis of LOT and cortical development, we consistently observed tdTomato labeling in the olfactory epithelium (OE) and vomeronasal organ (VNO) of all mice examined (Fig. 4A_1_-F_1_). Both regions contain sensory neurons whose development occurs concurrently with that of mitral cells (Marin-Padilla and Amieva, 1989; Shay et al., 2008; Rodriguez-Gil et al., 2015; Liberia et al., 2025). Moreover, CR expression has been proposed as a marker of intermediate stages of OSNs maturation in both the OE and VNO (Kishimoto et al., 1993; Wei et al., 2013). In the Calb2-CreERT2 × Ai14 mice, CR expression was first detected at E13 in the dorsal OE (Fig. 4B_1_, OSNs insets), and increased progressively at subsequent developmental stages (Fig. 4C_1_, D_1_, E_1_, F_1_, OSNs insets). In the VNO, the first evidence of CR expression in sensory neurons was at E16 (Fig. 4E_1_), from where it presumably will continue to rise until labeling the axons from the ON entering the accessory olfactory bulb (AOB) (Fig. 3C).

### Ultrastructural analysis of the LOT development and synaptogenesis in piriform cortex

Our analysis using the Calb2-CreERT2 × Ai14 mouse model provides a detailed characterization of the spatial organization and developmental dynamics of LOT axons. However, changes in axonal distribution and organization at the light-microscopic level do not necessarily reflect the ultrastructural maturation of the tract. To further characterize this process, we performed electron microscopy (EM), which enabled us to directly quantify axonal profiles and synaptic contacts within the LOT. First, we studied the density (number of axons/mm^2^) of LOT axons in APC at embryonic and postnatal ages to account for the incoming axons and maturation across development (Fig. 5), and then we analyzed synaptogenesis in layer Ia of PC (Fig. 6), the region where most LOT axons make synapses with pyramidal cells from layers II and III (Bekkers and Suzuki, 2013; Martin-Lopez et al., 2019a; Shepherd et al., 2021).

**Figure 6.**
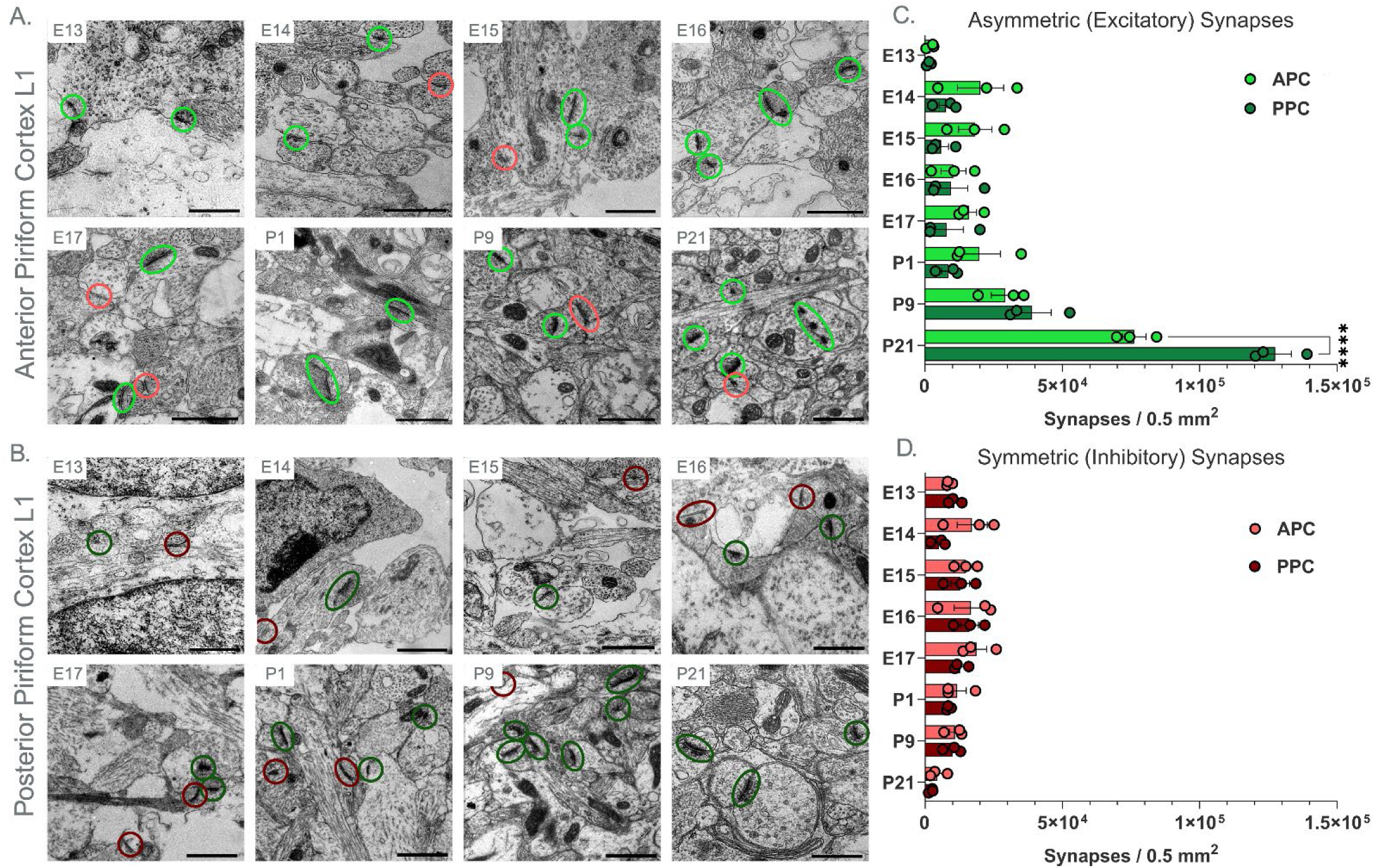
EM shows synaptogenesis across embryonic and early postnatal development in Layer Ia of PC. **(A)** Representative EM images of APC L1 selected from each age E13–E17, P1, P9, and P21. Light green circles represent asymmetric (excitatory) synapses. Light red circles represent symmetric (inhibitory) synapses. **(B)** Representative EM images of PPC L1 selected from each age E13–E17, P1, P9, and P21. Dark green circles represent asymmetric (excitatory) synapses. Dark red circles represent symmetric (inhibitory) synapses. **(C)** Quantification of asymmetric (excitatory) synapses per 0.5mm^2^ throughout development. Two-way ANOVA with Tukey’s multiple comparisons showing statistically significant differences between anterior and posterior PC at P21. **(D)** Quantification of symmetric (inhibitory) synapses per 0.5mm^2^ throughout development. Two-way ANOVA with Tukey’s multiple comparisons. Statistical significance: *p < 0.05. Scale bars: All 1μm across ages.

Regarding the LOT axons, we found that at E13 their morphology was very primitive with large accumulation of organelles (mitochondria) and prominent cytoplasm with vacuoles (Fig. 5-E13, asterisks), typical of axons that undergo active growth and extension (Tennyson, 1970), and their diameter was variable. At this age, the density of LOT axons was significantly lower when compared to other embryonic ages but not postnatally at P9 and P21, which we attributed to the thickening of axons during myelination (Fig. 5A-E13). After E14 the LOT was characterized by a sharp increase in the axon density showing more mature and regular morphologies with smaller diameters (Fig. 5A-E14). Throughout the remainder of embryonic development, the density of LOT axons remained relatively constant (Fig. 5A). Postnatally, starting at P9, myelination of LOT axons was evident and continued to P21, with a thickening of their diameters that resulted in lower density of LOT axons (Fig. 5A.). Interestingly, axon collaterals were observed at all embryonic and early postnatal stages examined (Fig. 5, blue arrowheads). Importantly, these measurements reflect axon density within the LOT and did not account for the overall increase in LOT size resulting from the progressive maturation of mitral/tufted cells and the addition of their axons to the tract. This progressive expansion of the OB-LOT is a well-established feature of olfactory system development and occurs during both embryonic and postnatal stages (Lopez-Mascaraque et al., 2005; Walz et al., 2006; Sarma et al., 2011; Collins et al., 2018).

Finally, we investigated synaptogenesis in layer Ia of PC during the embryonic and early postnatal development. Our previous IHC analysis for synaptophysin suggested that synapse formation begins at E13; therefore, we selected this age as the starting point for our EM analysis. To determine whether synaptogenesis differed along the rostral–caudal axis as LOT axons extended into the PC, we quantified the densities of asymmetric (excitatory) and symmetric (inhibitory) synapses in the anterior (APC) and posterior (PPC) PC based on their ultrastructural features (Fig. 6A, B). Because some LOT axons had already reached the PPC by E13 (Fig. 4), we were able to identify the first synapses in both the APC and PPC at this age (Fig. 6A–D), consistent with previous observations (Derer et al., 1977). Throughout the remainder of embryonic development, the densities of both asymmetric and symmetric synapses remained relatively low, with no significant differences between ages or regions (Fig. 6A–D). These findings suggested that synaptic development proceeded similarly in the APC and PPC, despite their different distances from the OB. By P9, we detected an increase in the density of asymmetric synapses in both regions, which became statistically significant by P21, indicating substantial postnatal maturation of excitatory connectivity in the PC. This increase coincides temporally with the pronounced period of myelination reported in the PC between P10 and P21 (Moriizumi et al., 1995). In contrast, symmetric synapse density remained relatively stable throughout development (Fig. 6B, D). Collectively, these findings identify E13 as the earliest age at which synapses were detected in layer Ia of the PC, with both excitatory and inhibitory synapse densities remaining steady over the entire embryonic development. Postnatally, asymmetric synapses showed a rapid increase especially by P21, correlating with an increase in myelination. The similar developmental synaptic profiles observed in APC and PPC suggested that synaptic maturation proceeds largely in parallel along the anterior–posterior axis, rather than being tightly coupled to the timing of LOT axon arrival in each region.

## Discussion

This study represents a novel approach to investigating the development of the lateral olfactory tract (LOT) in mice using calretinin (CR) as a proxy marker for olfactory bulb (OB) projection neurons. We aimed to address existing discrepancies in the literature regarding the sequential steps underlying LOT axonal elongation and maturation, particularly during early development. To this end, we combined immunohistochemistry (IHC) with Calb2-CreERT2 × Ai14 transgenic mice that express a fluorophore under the regulation of the CR promoter to track mitral/tufted maturation and axonal growth. At adult stages we found that most mitral/tufted cells expressed CR, albeit at lower levels than CR^+^ interneurons. This difference in mitral/tufted cell labeling was especially pronounced in dorsal OB and likely accounted for the reduced CR labeling observed in the ventral portion of the LOT within the piriform cortex (PC), which we termed the low-CR-labeling region. Embryonically, we detected early CR expression in a subpopulation of mitral cells by E12, whose axons had already reached posterior regions of PC and amygdala. Collectively, our findings provide a detailed timeline of LOT development in the PC and amygdala and establish CR labeling as a useful approach for investigating the development of olfactory circuitry and the axonal projections of OB projection neurons.

The use of CR as a marker for the LOT is not new, as several of our previous studies used CR immunolabeling to distinguish the laminar organization of the PC and LOT (Sarma et al., 2011; Martin-Lopez et al., 2019a; Martin-Lopez et al., 2019b). These findings prompted us to consider CR as a potential marker for mitral/tufted cells and, consequently, as a tool for investigating LOT development. We therefore began by characterizing CR expression in both the OB and PC, to have a better sense of how this molecule distributes in projection neurons. In the OB, we found that mitral/tufted cells expressed CR across the circumference of the OB (Figs. 1 and 2). This finding contrasts with previous reports describing either an absence of CR expression in mitral cells or its restriction to cells located in the lateral OB (Jacobowitz and Winsky, 1991; Resibois and Rogers, 1992; Wouterlood and Härtig, 1995). However, we did find lower CR expression in projection neurons located in the dorsal OB, which may account for the low-CR-labeling region identified in the LOT. These findings suggested a potentially unexpected topographical relationship in which axons originating from the dorsal OB contribute to the ventral portion of the LOT. If this topology is confirmed, it could challenge the current view of LOT organization, which proposes that axons originating from the dorsal OB travel predominantly through the dorsal LOT (Shepherd and Haberly, 1970; Hintiryan et al., 2012).

We next sought to establish a method for using CR to label and track OB projection neuron axons during development. Our initial analysis revealed that CR IHC did not reliably visualize individual axons. Because CR antibodies detect the protein itself, we hypothesized that the diffuse labeling observed in the LOT reflected the low abundance of CR in mitral/tufted cell axons, resulting in weak but detectable immunoreactivity. Similar labeling patterns have been reported for other proteins, including Reln and BRN2 (Alcantara et al., 1998; Ramos-Moreno et al., 2006; Brunjes and Osterberg, 2015). To overcome this limitation, we used the Calb2-CreERT2 × Ai14 mouse model, in which a reporter fluorophore, tdTomato, is expressed in CR^+^ cells instead of the protein itself. As expected, tissue sections from these mice revealed mitral/tufted cell axons and, consequently, the LOT. However, the low-CR-labeling region observed by IHC was absent in the PC. This result was expected because tdTomato expression does not depend on the level of CR protein, but rather on CreERT2-mediated recombination at the ROSA26 locus. Following 4OHTx administration, CreERT2 is activated and excises the STOP cassette, thereby placing tdTomato expression under the control of the ubiquitous CAG promoter (see Materials and Methods for details). Consequently, cells with different levels of CR expression can undergo recombination and subsequently express tdTomato at comparable levels. Consistent with this mechanism, tdTomato labeling was relatively uniform throughout the LOT and layer Ia of the PC (Fig. 3b_3_). An additional observation was the stronger CR immunolabeling in layer Ia of the PC compared with the LOT itself, a difference that was also absent in the transgenic mouse model. We speculate that this increased immunoreactivity in layer Ia may reflect the accumulation or extracellular release of CR at axon terminals, potentially through a mechanism similar to the secretion of Reln proposed by Martinez-Cerdeño et al. (2003). However, further investigation will be required to determine whether CR is released from mitral/tufted cell axon terminals. Taken together, these findings support our interpretation that the low-CR-labeling region detected by IHC reflects reduced CR abundance in axons traversing the ventral LOT, consistent with the lower CR expression observed in mitral/tufted cells located in the dorsal OB.

We subsequently studied the development of the LOT, including the differentiation and maturation of mitral/tufted cells in Calb2-CreERT2 × Ai14 mice following 4OHTx administration at E11. For the first time we showed that some pioneer mitral cells that expressed CR were already differentiated with non-arborized apical dendrites and extending axons by E12 (Fig. 4A1, MC inset). We found these axons reached posterior regions of the brain, in the prospective PPC and AMY, which occurred earlier than previously thought (Hinds, 1972; Sato et al., 1998; Walz et al., 2006). Nevertheless, our analysis of the LOT formation revealed unique developmental steps such as the innervation first of the amygdala compared to PPC, the latter occurring fundamentally after E15. All these processes correlated with the establishment of the first synapses that remained steady and in low densities across the entire embryonic development, reinforcing the idea that the majority of the activity in the olfactory system occurs postnatally (Moriizumi et al., 1995; Bartolomei and Greer, 1998; Martin-Lopez et al., 2018). The density of embryonic synapses is likely to be related with the emergence of spontaneous activity in PC during the refinement of circuits before spiking after birth (Barnett et al., 2014; Wang et al., 2024).

In summary, our work provides a comprehensive analysis of the development and molecular organization of the LOT : (1) OB mitral/tufted cells express CR, although at lower levels than interneurons; (2) the expression of CR by OB projection neurons is decreased in the dorsal region; (3) the ventral portion of the LOT within the PC contains a low-CR-labeling region that likely reflects the contribution of axons originating from the dorsal OB; (4) Calb2-CreERT2 × Ai14 mice provide a suitable model for labeling the somas, axons, and dendrites of CR^+^ mitral and tufted cells, enabling detailed analysis of LOT development; (5) pioneer CR^+^ mitral cells are detectable as early as E12 and extend axons toward posterior regions of the brain; (6) early LOT axons reach first the amygdala between E12-E14, followed by the PPC at approximately E15; (7) the axonal density of the LOT remains relatively stable throughout embryonic development, whereas postnatally decreases as axons increase in diameter and become progressively myelinated; (8) synaptogenesis in layer Ia of the PC does not follow an anterior-to-posterior gradient corresponding to the sequential arrival of LOT axons; and (9) layer Ia undergoes a marked increase in excitatory (asymmetric) synapses at P9, accompanied by a decline in inhibitory (symmetric) synapses.

## Acknowledgements

The authors express their appreciation to Dr. Jessica Cardin for advice on AAV injections into the transverse sinus.

## Credit author statement

Conceptualization: E.M-L., C.A.G.; Formal analysis: E.M-L., H.L., N.J.S., C.A.G.; Funding acquisition: C.A.G., Investigation: E.M-L., H.L., N.J.S., N.B., J.M.E-G., B.B., C.K., C.A.G.;

## Methodology

E.M-L., H.L., N.J.S., N.B., J.M.E-G., B.B., C.K.; Project administration: C.A.G.;

## Resources

C.A.G.; Supervision: E.M-L., C.A.G.; Validation: E.M-L., H.L., N.J.S., C.A.G.; Visualization: E.M-L., H.L., N.J.S., N.B., J.M.E-G., C.A.G.; Writing – original draft: E.M-L., C.A.G.; Writing – review & editing: E.M-L., H.L., C.A.G.;

## Supplementary Material

**Figure S1.**
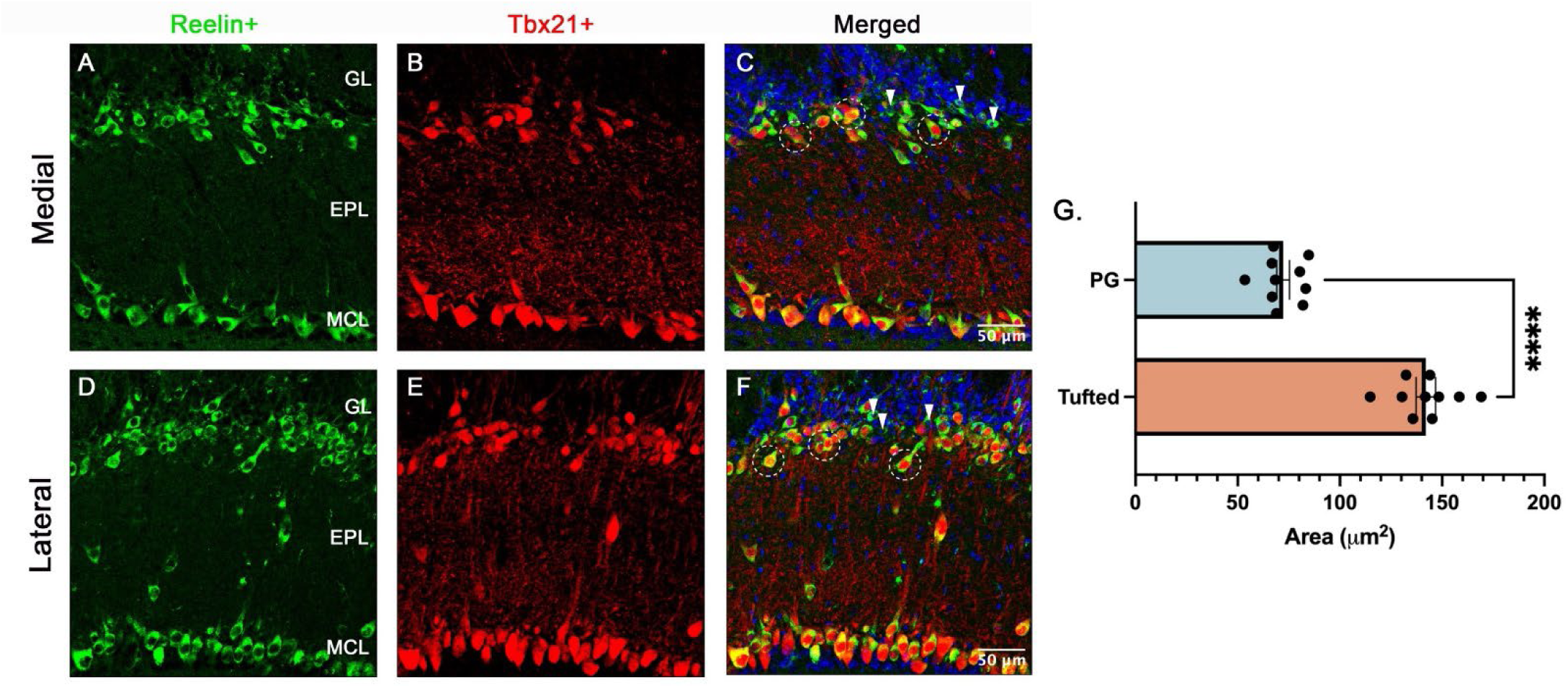
Tufted cells and PGNs can be distinguished in OB based on mean soma area. All figures show Reln (Mouse antibody) in green and Tbx21 (Rabbit antibody) in red. Medial OB: **(A)** Single-channel Reln, **(B)** Single-channel Tbx21, **(C)** Reln and Tbx21 colocalization. Lateral OB: **(D)** Single-channel Reln, **(E)** Single-channel Tbx21, **(F)** Reln and Tbx21 colocalization. **(G)** Quantification of mean soma area between tufted cells and PGNs. Overall, Tufted cells exhibit significantly greater soma area. Unpaired t-test. Statistical significance: ****p < 0.0001. Scale bars: 50μm in (C, F).

**Figure S2.**
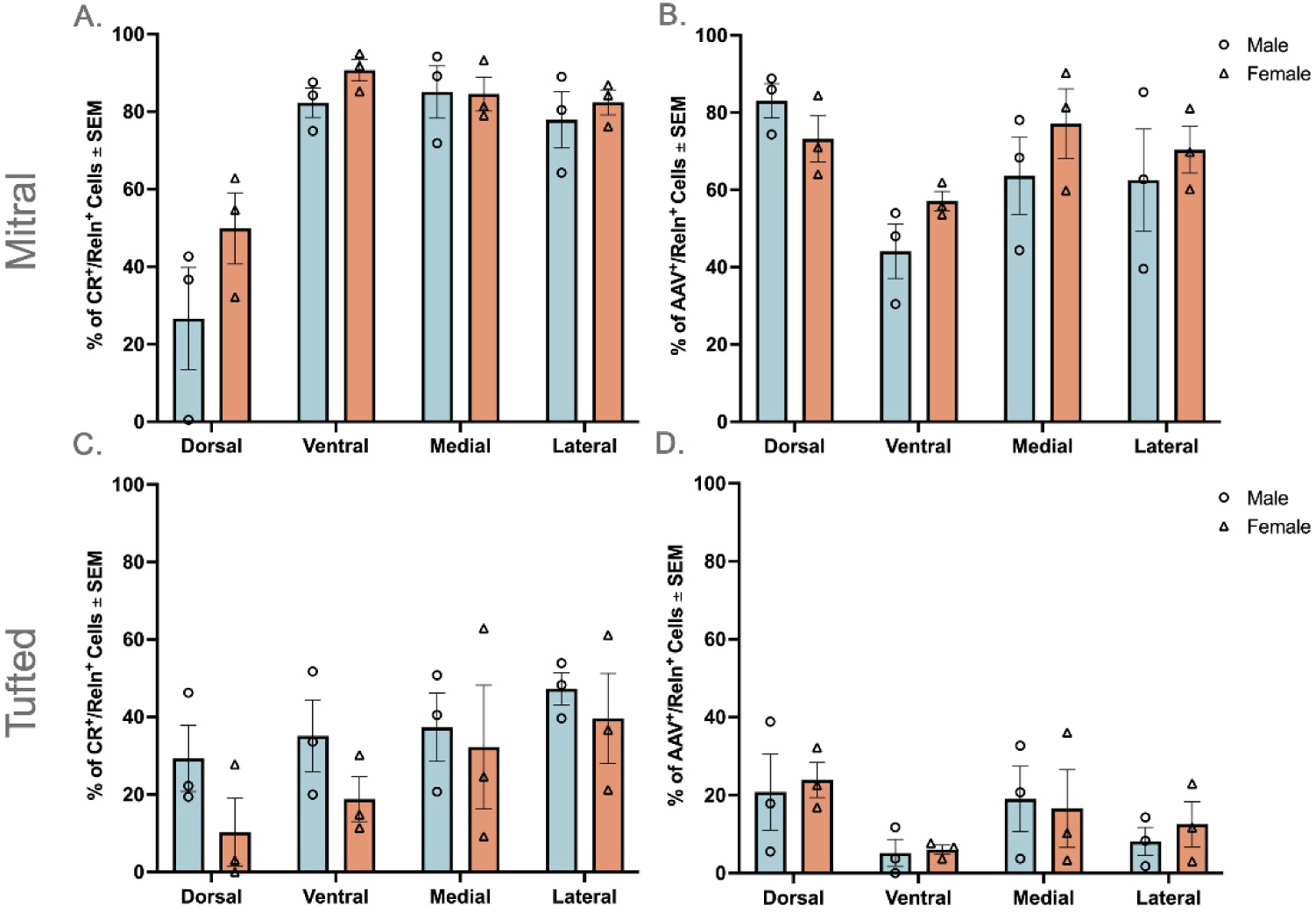
No statistical differences between sexes were observed for both CR and AAV groups. **(A, B)** Quantifications involving mitral cells (n = 6; 3 males, 3 females): **(A)** Percentage of CR^+^/Reln^+^ cells across regions split by sex (multiple unpaired t-tests using Holm-Šídák method) and **(B)** Percentage of AAV^+^/Reln^+^ cells across regions split by sex (multiple unpaired t-tests using Holm-Šídák method). **(C, D)** Quantifications involving tufted cells (n = 6; 3 males, 3 females): **(C)** Percentage of CR^+^/Reln^+^ cells across regions split by sex (multiple unpaired t-tests using Holm-Šídák method) and **(D)** Percentage of AAV^+^/Reln^+^ cells across regions split by sex (multiple unpaired t-tests using Holm-Šídák method). No statistical significance was observed.

